# bpRNA: Large-scale Automated Annotation and Analysis of RNA Secondary Structure

**DOI:** 10.1101/271759

**Authors:** Padideh Danaee, Mason Rouches, Michelle Wiley, Dezhong Deng, Liang Huang, David Hendrix

## Abstract

While RNA secondary structure prediction from sequence data has made remarkable progress, there is a need for improved strategies for annotating the features of RNA secondary structures. Here we present bpRNA, a novel annotation tool capable of parsing RNA structures, including complex pseudoknot-containing RNAs, to yield an objective, precise, compact, unambiguous, easily-interpretable description of all loops, stems, and pseudoknots, along with the positions, sequence, and flanking base pairs of each such structural feature. We also introduce several new informative representations of RNA structure types to improve structure visualization and interpretation. We have further used bpRNA to generate a web-accessible meta-database, “bpRNA-1m”, of over 100,000 single-molecule, known secondary structures; this is both more fully and accurately annotated and over 20-times larger than existing databases. We use a subset of the database with highly similar (≥90% identical) sequences filtered out to report on statistical trends in sequence, flanking base pairs, and length. Both the bpRNA method and the bpRNA-1m database will be valuable resources both for specific analysis of individual RNA molecules and large-scale analyses such as are useful for updating RNA energy parameters for computational thermodynamic predictions, improving machine learning models for structure prediction, and for benchmarking structure-prediction algorithms.

## INTRODUCTION

Ribonucleic acid (RNA) is a type of macromolecule that is essential for all life, with functions including molecular scaffolding, gene regulation, and encoding proteins. The secondary structures and base-pairing interactions of RNAs reveal information about their functions(1–4). While RNA structure prediction has made tremendous improvements in the past decades, there are several limitations in available resources for researchers. While over 100,000 known RNA structures exist in various databases,the most detailed meta-database, RNA STRAND v2.0, contains less than 5,000 entries, and has not been updated in a decade. Moreover, even with base pairing data, the structural features present can be rather complex and there has not yet been a fully successful general approach presented to systematically resolve the structural topology and identify all structural features given the base pairing information. This limitation is part of the reason that most source databases do not provide dot-bracket sequences for their structures. Therefore, there is a need for reliable tools that identify and annotate structural features from RNA base pairing data.

We present “bpRNA”, a fast, easy-to-use program that parses base pair data into detailed structure “maps” providing relevant contextual data for stems, internal loops, bulges, multi-branched loops (multiloops), external loops, hairpin loops, and pseudoknots. Previous work to parse RNA structural topology from base pairs do not handle pseudoknots(5). bpRNA outputs new file formats (both high-level and detailed-level) for RNA secondary structures that provide information to help understand the structure. bpRNA has accurately generated the dot-bracket sequence for all structures, including the complex structures with pseudoknots.

The prediction of RNA secondary structure is based on thermodynamic model parameters that are calculated from available data of known structures(6–8). Likewise, the study of RNA secondary structure creates a need for comprehensive meta-databases, the analysis of which could enable updated RNA thermodynamic parameters and prediction tools. The detailed structural annotations generated by bpRNA provide information needed to build a rich database of great use to the RNA research community. While databases of 3D structures exist(9–11), they don’t serve the same needs as secondary structure databases. There have been many attempts at creating RNA secondary structure databases and meta-databases(12–14), all of the meta-databases except RNA STRAND v2.0(13) are no longer available or have not been updated. To meet this need, we have built a detailed meta-database, “bpRNA-1m”, consisting of over 102,318 single molecule (1m) RNA secondary structures extracted from seven different sources, and analyzed by bpRNA. These data, including the structure annotations provided by bpRNA, represent the largest and most detailed RNA secondary structure meta-database created to date and will be expanded as more data become available. This comprehensive meta-database can be used in machine learning applications, benchmarking studies, or can be filtered as desired for other RNA structure research.

## MATERIAL AND METHODS

### RNA Secondary Structure Types

We use the term “stem”, as previously defined(13), to refer to a region of uninterrupted base pairs, with no intervening loops or bulges (Figure 1A). We label the two paired sequences of a stem as 5’ or 3’ depending on their order in the RNA sequence. A hairpin loop is an unpaired sequence with both ends meeting at the two strands of a stem region. The direction of the hairpin loop sequence also defines the nucleotides in the closing base pair and mismatch pair as being 5’ or 3’ (Figure 1B). An internal loop is defined as two unpaired strands flanked by closing base pairs on both sides, which are labeled as 5’ vs 3’ based on which is more 5’ in the RNA sequence (Figure 1C). The closing base pair 5’ of the 5’ strand is labeled as the 5’ closing pair, and the closing pair that is 3’ of the 5’ strand is the 3’ pair. A bulge can be considered as a special case of the internal loop where one of the strands is of length zero (Figure 1D). Multi-branch loops (multiloops) consist of a cycle of more than two unpaired strands, connected by stems (Figure 1E). External loops are similar to multiloops, but are not connected in a cycle. Dangling ends are unpaired strands at the beginning and end of the RNA sequence.

**Figure 1:**
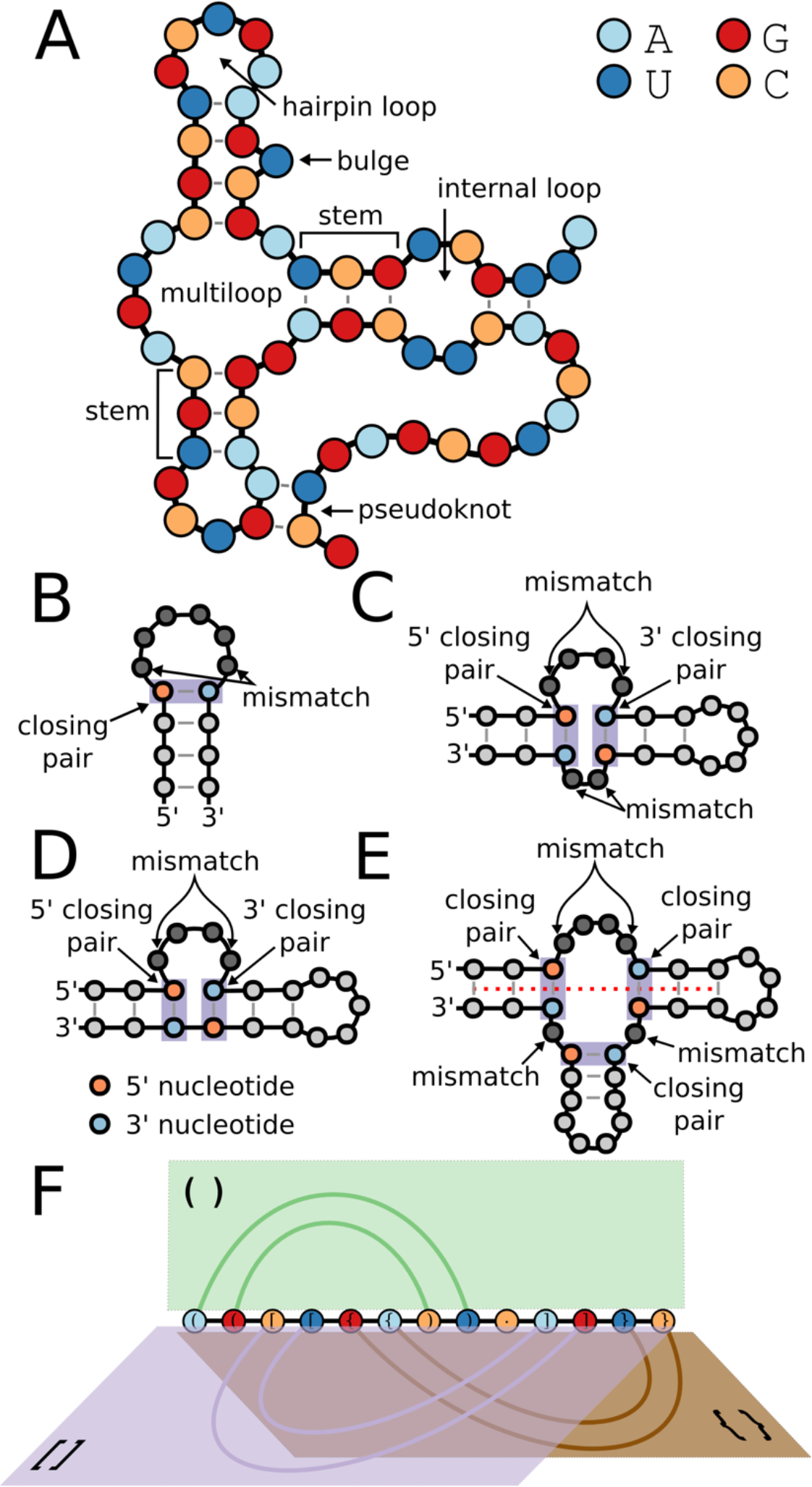
RNA Structure Types. **A.** cartoon schematic of RNA structure types. **B.** Hairpins have one closing pair and one mismatch pair with nucleotides defined by ordering from 5’ to 3’. **C.** Internal loops have two closing base pairs and two mismatch pairs each defined by ordering from 5’ to 3’ relative to the 5’ internal loop strand. The nucleotides of the closing pairs are defined as 5’ or 3’ based on their positions relative to the loop sequence. **D.** Bulges have one loop strand, but have two closing base pairs and two mismatch pairs defined 5’ to 3’. **E.** Multiloops have a closing pair for each branch. The nucleotides of the closing pairs are defined as 5’ or 3’ based on their positions relative to the loop sequence. Red dashed line represents the common axis of coaxially stacked stems. **F.** A depiction of RNA page number, which can be viewed as separate half-planes containing edges corresponding to base pairs. Each symbol type corresponds to a separate page, and edges within a page are nested.

Pseudoknots (PKs) are characterized by base-paired positions *i*, *j* and (*i*’, *j*’) that satisfy the PK-ordering, which is defined as either *i* < *i*’ < *j* < *j*’ or *i*’ < *i* < *j*’ < *j*. For a secondary structure, PK base pairs are annotated as the minimal set that result in a PK-free structure when removed(5,13,15–17). While representations of PK-containing RNA structures are not planar, the “book embedding”, or the number of distinct half-planes with a common boundary line (the RNA strand) can describe the RNA structure(18). The number of half-planes needed to represent the structure is called the “page number”, and a book embedding for an RNA structure that has a lower page number is preferred because it provides a more compact representation. Figure 1F depicts the pages for an RNA structure with a page number of 3.

### Segment Graph representation

We have defined the “segment” and “segment graph” to assist in parsing RNA secondary structures and for visualization of structures. A segment is a region consisting of two strands of duplexed RNA that can contain bulges or internal loops. The difference between a stem and a segment is that segments can contain unpaired bases. When a base pair at positions (*i*, *j*) is part of a segment, then if the next paired nucleotide 5’ of *i* is paired to the previous paired nucleotide 3’ of *j* then this base pair is also part of the segment (See Supplementary Methods). As an illustration of this idea, Figure 2A presents the structure of a ribozyme that contains 8 color-coded segments numbered 5’ to 3’. This definition allows us to parse a structure into segments in linear time (“IdentifySegments” Algorithm in Supplementary Methods). The segment concept has some similarity to “bands”, which is loosely defined as “a pseudoknotted stem, which may contain internal loops or multi loops”(19), except segments apply more generally than pseudoknots, and do not contain multiloops. Pseudoknots (PKs) can be segments as well, such as segments 1 and 5 in Figure 2A-B, but the concept generally applies to any paired region.

**Figure 2:**
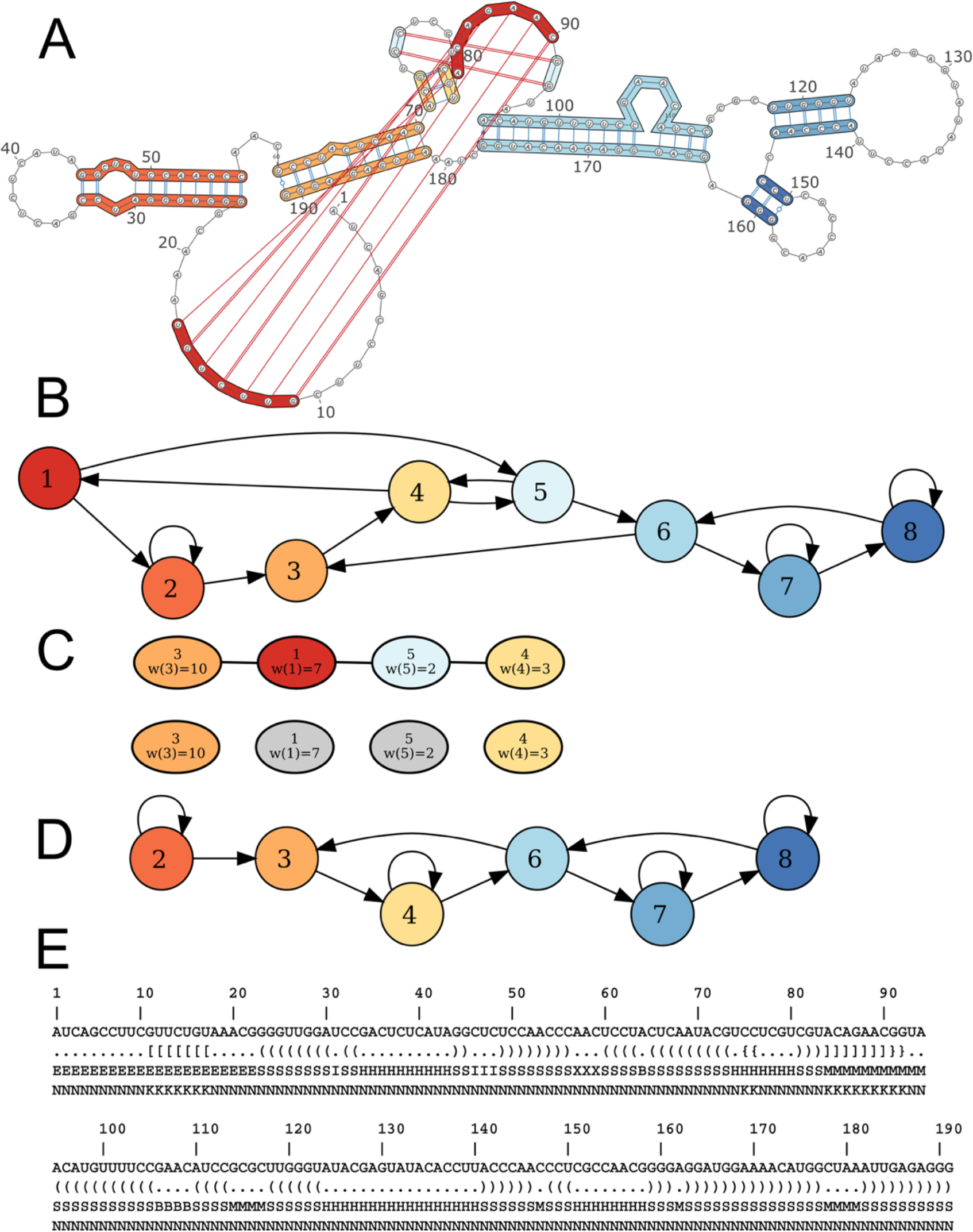
Segment Graph example. **A.** Secondary structure of the *Anopholes gambia* drz-agam-2-2 ribozyme. **B.** The segments are the vertices of the segment graph and ordered from 5’ to 3’, and directed edges are defined by unpaired strands connecting segments. **C.** Segments with base pairs crossing other segments comprise the PK-graph. A maximally weighted independent set is selected by dynamic programming, with the remaining segments defined as pseudoknots. **D.** The pseudoknot-free segment graph is created after remove PK base pairs and allows easy annotation of loops. **E.** The structure array enhances bpRNA’s multi-bracket dot-bracket sequence by labeling each positions structure type. Strands participating in pseudoknots are labeled in the structure array by their loop-type in the structure resulting from the removal of PKs.

The upshot of the segment representation is we can create a “segment graph”, which provides a compact representation of each structure (Figure 2B). Others have defined graph representations of RNA structures, such as “RNA As Graph”(20,21); however, the problem is these representations use stems as the edges of an undirected graph, making this extraordinarily complex for typical long noncoding RNAs, which can contain hundreds of stems or more. Moreover, examples from biology such as microRNAs show that many structures can preserve their functionality even when including bulges and internal loops(22). These examples suggest a value in a more coarsely-defined secondary structure graph concept.

For any structure, we can define a directed multigraph *G* = (*V*,*E*) such that the vertices *V* of the graph are the segments, the directed edges *E* correspond to unpaired strands, in the 5’ to 3’ direction, connecting them. Two segments can have an edge even when there is no intervening unpaired nucleotide (only a backbone). Each vertex of a segment graph can have at most two outgoing and two ingoing edges. Only the first and last segment can have less than two ingoing and outgoing edges. Hairpin stem-loops are easily identified as segments with self-edges, which count as one outgoing and one ingoing edge.

Pseudoknots (PKs) have been identified previously as the minimum set of base pairs that, when removed, produce a pseudoknot-free structure(5,15–17), and algorithms have been developed for optimal selection of these base pairs(23). We use the segment concept to identify this minimal set of base pairs. All pseudoknot base pairs are part of a segment, and these pseudoknot segments (PK-segments) can be easily identified; if one base pair of a segment satisfies the PK-ordering with a base pair in another segment, then all base pairs in this segment satisfy the PK-ordering with all base pairs in the other segment (See proof in Supplementary Methods). Once PK-segments have been identified, a weighted, undirected graph called a PK-segment graph can be created such that the PK-segments correspond to vertices and edges connect them when they satisfy the PK-ordering with each other (Figure 2C). We assign a weight to each vertex, with the value of the number of base pairs for the PK-segment. From this graph, we next identify the maximum weighted independent subset (MWIS), leaving a minimal subset of segments whose removal leaves the secondary structure free of pseudoknots (Figure 2C). We created an exact algorithm, “MaxPKFreePairs” to selecting the MWIS using a Nussinov-style(24) dynamic programming approach similar to defined previously(5), as well as a heuristic algorithm “PK_Detection” for dealing with ties (Supplementary Methods). We found that both methods produce the same solution to identifying the minimum subset of base pairs needed to produce a PK-free structure. These segments are then annotated as pseudoknots, and can be excised to produce a pseudoknot-free (PK-free) structure and PK-free segment graph (Figure 2D). The PK-free structure is equivalent to the page number=1 structure. The full algorithm for this approach is presented in the Supplementary Methods.

The PK-free segment graph enables facile identification of structure types. Hamiltonian cycles in the PK-free segment graph correspond to multiloops (Figure 2D). Interior loops and bulges can be identified as unpaired bases within segments. Pseudoknots are not discarded, but rather we annotate pseudoknots by the type of loops they connect in the corresponding PK-free structure. For instance, if a PK consists of base pairs connecting what would otherwise be a multiloop branch and a bulge, we label the PK as “M-B”.

The bpRNA code is written in perl and requires the Graph perl module. Several additional scripts for analysis are included. The source code is available at http://github.com/hendrixlab/bpRNA

### Reference Databases

The seven databases that comprise the bpRNA-1m meta-database include Comparative RNA Web (CRW) (25), tmRNA database(26), tRNAdb (27), Signal Recognition Particle (SRP) database(28), RNase P database(29), tRNAdb 2009 database(30), and RCSB Protein Data Bank (PDB) (31), and all families from RFAM 12.2(32). Moreover, to reduce duplication for further analysis, we created a subset called bpRNA-1m(90), where we removed sequences with greater than 90% sequence similarity when there is at least 70% alignment coverage(33). The bpRNA-1m database currently has 102,318 RNA structures and the bpRNA-1m(90) subset consists of 28,370 structures. For comparison, the RNA STRAND v2.0 database has 4,666 structures, with fewer than 2,000 structures when similarly filtered.

The Comparative RNA Web (CRW) site contains RNA sequences and secondary structures obtained from comparative sequence analysis. There are 55,600 records extracted from this reference through the mass data retrieval option. For each RNA extracted from this source, we retrieved phylogenetic lineage, organism name, and RNA type. The tmRNA Database provides structures of transfer messenger RNAs (tmRNAs), which are bacterial RNAs with both tRNA- and mRNA-like functions. The base pair information for all 728 RNAs from this source was also determined using comparative sequence analysis. Single Recognition Particle Database (SRP) is a source for structures and functions of single recognition particle RNAs (SRP RNAs) along with phylogenetic lineage and organism names for each RNA(27). The tRNAdb 2009 database (formerly Sprinzl tRNA Database) encompasses all the structures and sequences from tRNA genes from three different university sources: Leipzig, Marburg, and Strasbourg(30). All 623 of these verified RNA structures were downloaded from this source along with their taxonomy and links to each individual reference. The RNase P Database (RNP) has sequences and secondary structures of Ribonuclease P type RNA of bacteria, archaea, and eukaryotes. All available taxonomy, organism name, and associated PubMed ID data were downloaded for the 466 entries in this database.

RCSB Protein Data Bank (PDB) contains structures of proteins and nucleic acids obtained using X-ray crystallography and NMR techniques. We download all 669 RNA structures (PDB files) consisting of one RNA molecule as of June 12 2017. We first parsed the 3D structures from PDB files with the June 2017 version of RNAview(34), and used custom perl scripts to convert to BPSEQ format. This conversion considers both canonical and non-canonical base pairs. The priority is on the positions with Watson-Crick and Wobble pairs. The Watson-Crick pairs are identified by the edge represented in RNAView output (+/+ for GC pairs and −/− for AU pairs), and wobble pairs are recognized when the edge is Watson-Crick/Watson-Crick and has the *cis* orientation with XXVII Saengers classification(34). Similarly, non-canonical pairs are extracted based on these three specifications(35).

The RNA Family Database (RFAM) V12.2 contains consensus structures derived from comparative sequence analysis of individual sequence family members of thousands of RNA families. For each sequence, we extracted the RNA type, validation technique and when available, the URL for the RNA family Wikipedia page. There are 2,495 families in RFAM V12.2 and 43,273 individual sequences. For each family, we projected consensus structures to individual sequences using multiple sequence alignments provide by RFAM and custom perl scripts. Base-paired positions in the consensus structure were mapped to individual sequences, while removing gaps in the alignment, as done in previous studies(13). We include information on the publication status in the database for users that want to exclude unpublished structures.

The relational database is implemented with MySQL (Version 15.1) on a CentOS GNU/Linux server (Supplementary Figure S1). For more detail on the database, see Supplementary Methods.

## RESULTS AND DISCUSSION

### The bpRNA approach

#### bpRNA Secondary Structure Decomposition and Representation

The input to bpRNA is a list of base pairs (BPSEQ file) for a given RNA secondary structure. First, the segments are identified as in Figure 2A,B. Next, a PK-graph is built, and the PK-segments are identified (Figure 2C). The PK-free segment graph, which enables multiloops and external loops to be easily identified, is built after the removal the base pairs in PK-segments (Figure 2D). Bulges and internal loops are identified as unpaired positions within the segments. After all loops are identified, the pseudoknots are annotated by the loops in the PK-free structure that they connect (See Methods). The output of bpRNA analysis are 1) a multi-bracket dot-bracket representation of the secondary structure, 2) a “structure array” sequence providing more detail to the dot-bracket, and 3) a “structure type” file. The content of these files is described in the following sections.

#### An Accurate Dot-bracket representation of RNA secondary structure

Dot-bracket format represents base pairs with paired parentheses, unpaired nucleotides with dots, and pseudoknots with other brackets (“[”,“{”,“<”…). While most of the databases that the data was derived from does not include a multi-bracketed dot-bracket representation when pseudoknots are present, bpRNA has successfully created one for every structure. Each dot-bracket representation we created is sufficient to re-create the BPSEQ file using our multi-stack approach to parse the dot-bracket structure. The efficiency of a dot-bracket sequence is described by the “page number”, which is the number of different symbol types used to represent the dot-bracket structure(36). Our dot-bracket consist of dots “.” for unpaired bases, matched parentheses indicate nested base pairs for page 1, square brackets for page 2, curly braces for page 3, angle brackets for page 4, and pairs of upper/lower alphabetical characters (Aa, Bb,…, Zz) for higher page numbers. Base pairs on the same page do not cross each other, i.e., each page is pseudoknot-free (Figure 1F). We were able to represent all structures with a page number less than or equal to 7, and 99.46% of the structures with a page number of 2 or less. For all 1,497 structures were bpRNA differs from RNA STRAND v2.0, bpRNA produced a lower page number lower page number, and thus a simpler dot-bracket sequence (Supplementary Figure S2). In some cases, RNA STRAND v2.0 had a page number as high as 30, requiring every letter of the alphabet to represent the pseudoknots of the structure, while bpRNA has a page number of 5.

#### The bpRNA “structure array”

We also created what we call the “structure array”, which is a series of single character identifiers for the structure types of each nucleotide in the sequence, providing another layer of annotation to supplement the dot-bracket (Figure 2E) and a high-level representation of the structure. In this representation, S=stem, H=hairpin loop, M=multi-loop, I=internal loop, B=bulge, X=external loop, and E=end. The next sequence labels nested or unpaired nucleotides with “N”, and nucleotides forming pseudoknots with “K”. This enables a compact representation and additional detail, making the dot-bracket more easily interpretable for researchers. This is particularly helpful for loop regions, which are only represented as a dot “.” in dot-bracket, and detailed by the type of loop with the structure array. Similarly, the structure array at pseudoknot positions indicates what loops result from the removal of the pseudoknots.

#### The bpRNA “structure type” file

We defined a new file format with each structural feature, relevant positions, and flanking base pairs, and sequences called a “structure type” file (Supplementary Figure S3). This file format goes beyond the dot-bracket and structure array because it has more detail such as positions and flanking base pairs, and is capable of representing features of length zero, such as zero-length multiloop branches. Each feature is numbered, and PK interactions are indicated for loops that contain them. Researchers can unambiguously explore a structure with this information, along with the dot-bracket, structure array, and VARNA 2D structure image.

#### bpRNA yields accurately annotated features

We found a number of differences with our feature extraction and other work. For bulges, we found 1,042 structures with differences in the identified number of bulges. For instance, Supplementary Figure S4 shows a structure for the tmRNA *List.wels._AF440351_1-321*. This is annotated as having 0 bulges in RNA STRAND v2.0, but it actually has 4 bulges and these are all correctly annotated by bpRNA. In Supplementary Figure S5, we label in the structure diagram from the RNA STRAND v2.0 database with the bulges identified by bpRNA, and provide location and sequence information for these bulges. For other structure types, we have a different classification system. For example, if a hairpin loop participates in a pseudoknot (e.g. a “kissing hairpin pseudoknot”), RNA STRAND v2.0 does not annotate it as a hairpin. In contrast, we still classify loops by the above definitions even when they contain nucleotides forming PK base pairs, but label them with the specific PK involved. Furthermore, we categorize PK base pairs by the loop sequences that they connect.

### The bpRNA-1m and bpRNA-1m(90) databases

The number of RNA structures extracted from each source is shown in Table 1 for both bpRNA-1m and bpRNA-1m(90). There are a relatively higher number of structures from CRW and RFAM database; however, around 92% of the CRW data are filtered running the CD-HIT-EST algorithm with the 90% similarity. In some cases, bpRNA detected errors in the source BPSEQ files used to build our meta-database: in cases where a nucleotide was paired to itself, the base pair was removed; in cases where a nucleotide was paired to two positions, one was removed. Overall, we found 30 such examples (Supplementary Table S1).

**Table 1.** The number of RNAs from each source is listed for both bpRNA-1m and bpRNA-1m(90)

|  | bpRNA-1m | bpRNA-1m(90) |
| --- | --- | --- |
| CRW | 55,600 | 4,368 |
| tmRNA | 728 | 339 |
| SRP | 959 | 352 |
| tRNAdb 2009 | 623 | 207 |
| RNP | 466 | 253 |
| RFAM | 43,273 | 22,521 |
| PDB | 669 | 330 |
| Total | 102,318 | 28,370 |

The complete 6 bpRNA-1m database is available through our interactive website at http://bpRNA.cgrb.oregonstate.edu

### Secondary Structure Feature Analysis

The output of bpRNA can help researchers understand RNA secondary structure, and enable large-scale structural analysis. As an example of the type of analysis that can be performed, we analyzed the resulting secondary structure annotations to identify enriched sequence and structural patterns in our database (Table 2). We performed this analysis on the bpRNA-1m(90) to reduce duplicated information. Table 3 shows the distribution of RNA types for bpRNA-1m and bpRNA-1m(90). We found several general trends in this large data set, which could be refined in future studies as more data become available, or with a more restricted subset.

**Table 2.** The number of each structures type for all RNA structures in bpRNA-1m and bpRNA-1m(90)

|  | bpRNA-1m | bpRNA-1m(90) |
| --- | --- | --- |
| Bulges | 517,672 | 82,061 |
| Hairpin Loops | 708,144 | 119,645 |
| Multiloops | 317,046 | 41,424 |
| Internal Loops | 538,670 | 93,435 |
| Pseudoknots | 57,686 | 7,164 |
| Exterior Loops | 229,468 | 67,059 |
| Stems | 2,075,928 | 335,877 |
| Segments | 1,019,586 | 160,381 |

**Table 3.** The number of common RNA types is listed for bpRNA-1m and bpRNA-1m(90)

| RNA Type | bpRNA-1m | bpRNA-1m(90) |
| --- | --- | --- |
| Transfer RNA | 35,622 | 3,383 |
| 16S Ribosomal RNA | 17,641 | 1,067 |
| 5S Ribosomal RNA | 4,577 | 607 |
| Signal Recognition Particle RNA | 1,603 | 388 |
| Ribonuclease P RNA | 1,425 | 605 |
| Transfer Messenger RNA | 1,361 | 449 |
| Group I Intron | 237 | 123 |
| 23S Ribosomal RNA | 191 | 72 |
| Hammerhead Ribozyme | 186 | 77 |
| Group II Intron | 131 | 101 |

#### Hairpin Loops

The most common loop-type found in RNA secondary structures are hairpin loops(37). For each hairpin loop, there is a closing base pair and unpaired region. The destabilizing energy of a hairpin loop can be determined from the type of the closing base pair, type of mismatch, and the length of the unpaired region (8,38). Using the bpRNA-1m(90), we found that tetraloops, hairpin loops of length four, are the most common (Figure 3A). While many previous studies focused hexaloops, loops of length 6 (8,39–41), we found that heptaloops, hairpin loops of length seven, are the second most frequent (Figure 3A). Hairpin loops of size less than 4 and greater than 7 occur much less frequently in bpRNA-1m(90).

**Figure 3:**
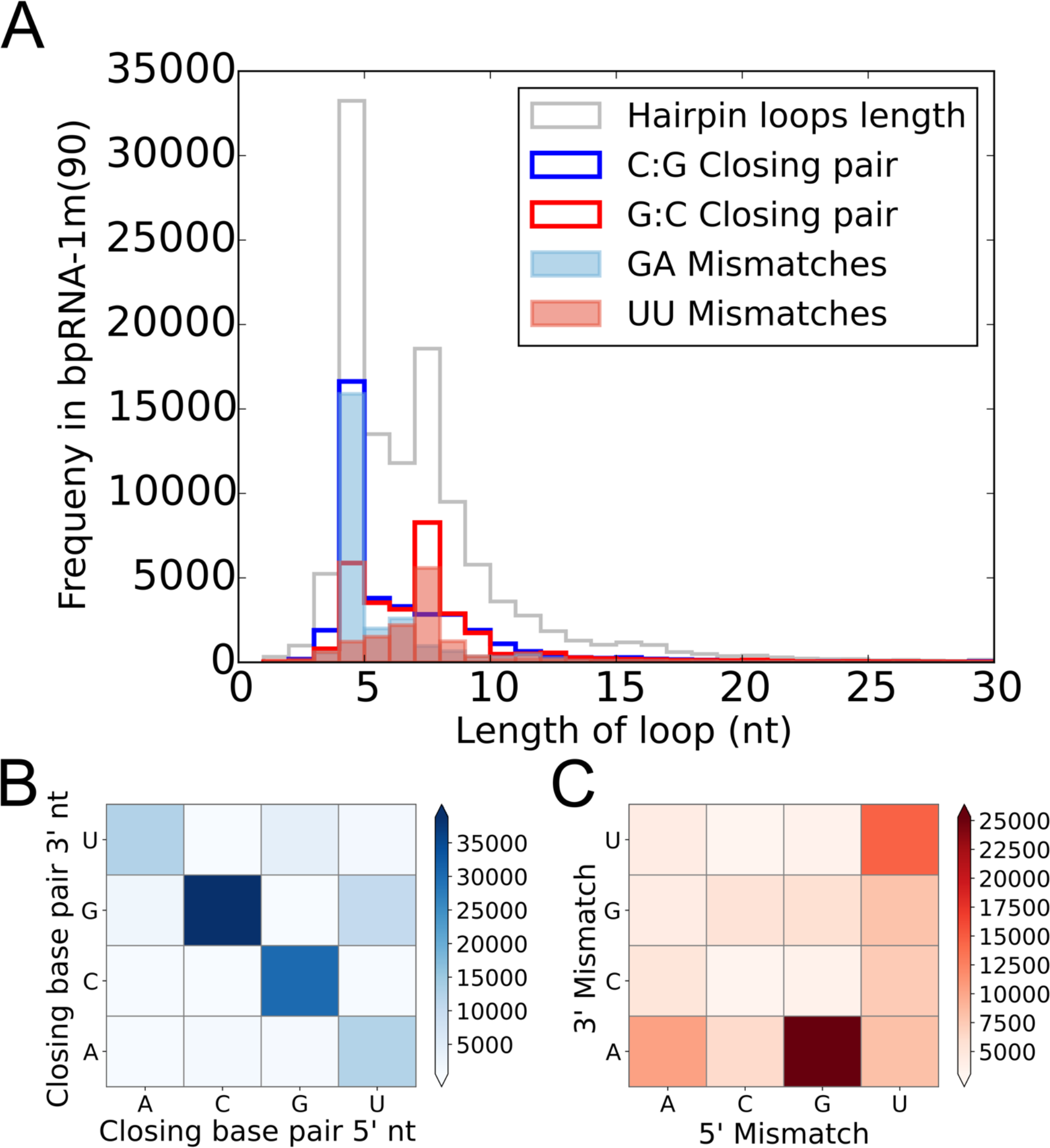
Hairpins in bpRNA-1m(90). **A.** The distribution of hairpin loop lengths in bpRNA-1m(90) has two primary peaks, overlapping the same peak for subsets defined by closing pairs. **B.** Heat map shows the frequency of nucleotides occurring in closing base pairs. **C.** Heat map shows the frequency of pairs of nucleotides occurring in hairpin mismatch pairs.

When considering all hairpin loops in bpRNA-1m(90), we found that C:G followed by G:C are the most common closing base pairs (Figure 3B), and GA mismatches are the most common overall (Figure 3C). The data suggest that tetraloops are significantly enriched with C:G closing base pairs, while heptaloops are enriched with G:C closing pairs. The tetraloops with C:G base pairs are mostly associated with GA mismatches, while heptaloops of size seven have the G:C base pair which is followed by UU mismatch. There are known frequent and stable patterns for tetraloops from various studies such as UNCG, GNRA, and CUUG, where N=A, C, G, or U and R=A or G (39,42,43). Previous work has compared the statistical frequency of secondary structural features to thermodynamic stability(44,45). Using Turner 2004 nearest neighbor model, we compared the destabilizing energy of the hairpin loop types to their frequency of occurrence in bpRNA-1m(90) (Figure 4A). As it is shown, the GNRA and UNCG patterns are highly abundant whereas CUUG was not as frequent in our set. Sequence LOGOs(46) for all tetraloop tokens and for the top 1% when sorted by type frequency is presented in Figure 4B. We also did the same energy calculation for heptaloops which is illustrated in Figure 4C along with sequence LOGOs in Figure 4D. Altogether in bpRNA-1m(90), the most common type for tetraloops is C(GAAA)G and for heptaloops the most common type is G(UUCGAAU)C (Figure 4). In other examples, there are loops that have low energy and a low frequency of occurrence. For example, G(GGUAAGC)U is probably rare because it is more stable for the GC mismatch to pair, forming a loop of length 5.

**Figure 4:**
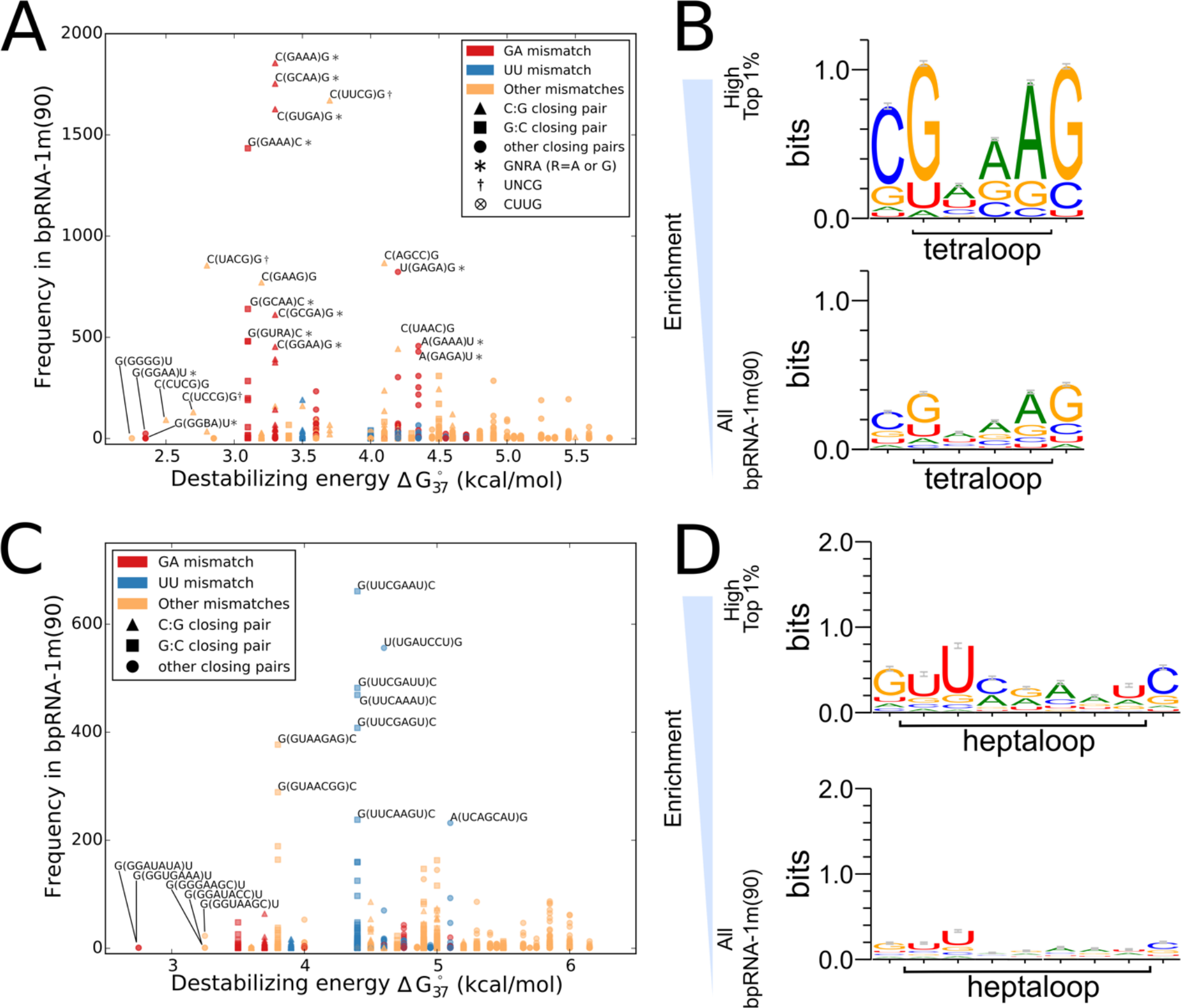
Tetraloops and Heptaloops. **A.** Scatterplot compares the frequency of tetraloop sequences to destabilizing energy. **B.** Sequence LOGOs demonstrate sequence biases in the most enriched tetraloops.**C.** Scatterplot compares the frequency of heptaloop sequences to destabilizing energy. **D.** Sequence LOGOs demonstrate the sequence biases in the most enriched heptaloops.

#### Internal Loops

Internal loops tend to be symmetric, because this creates a more stable structure(47). The internal loop frequency heat map (Figure 5A) demonstrates a tendency toward symmetric internal loops in bpRNA-1m(90), particularly when fewer than 4 nt. There are various factors in calculating the energy parameters of an internal loop such as first mismatch, closing base pairs, and the length of the 5’ and 3’ loop sequences(8). We found that while the 5’ closing base pair favors G:C, the 3’ closing base pair favors C:G (Figure 5B). Mismatch nucleotides, defined as the first and last nucleotide of the loop, are enriched for GA (Figure 5C). Moreover, we found that internal loops with GA mismatches were most likely to have a length of 3 (Supplementary Figure S6). We found that 5’ and 3’ internal loops had slightly different length distributions, with 5’ showing a greater propensity for length 3 (Figure 5D,E). The preference for C:G for the 3’ closing pair is especially true for 3’ internal loops longer than 3 nt (Figure 5E).

**Figure 5.**
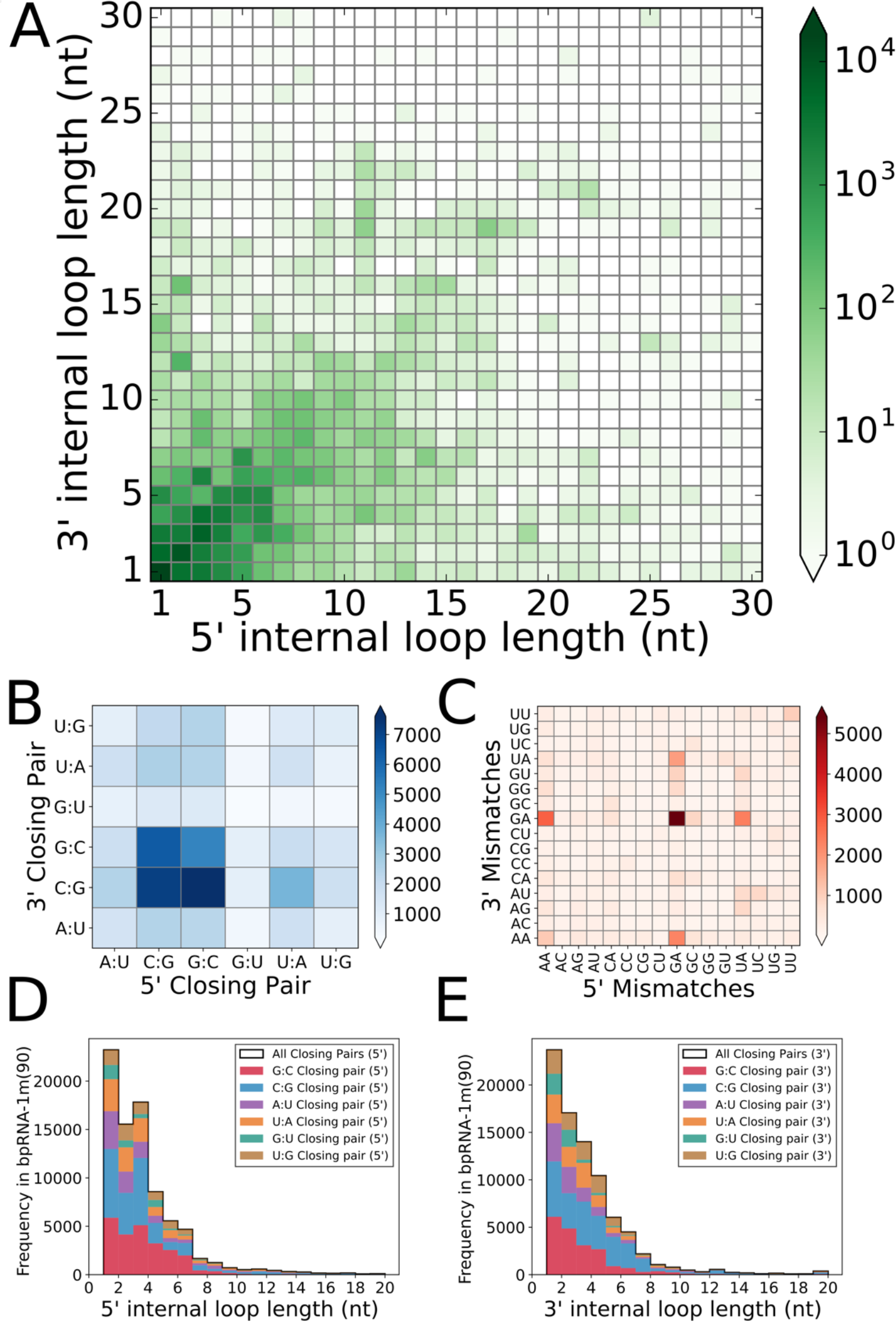
Internal loops. **A.** Heat map shows the frequency of internal loops based on 5’ and 3’ loop length.**B.** Heat map shows the frequency of base pairs occurring in 5’ and 3’ internal loop closing base pairs. **C.**Heat map shows the frequency of pairs of nucleotides occurring in 5’ and 3’ internal loop mismatch pairs.**D.** Stacked histograms of 5’ internal loop lengths when organized by the 5’ closing base pair. **E.** Stacked histograms of the 3’ internal loop lengths when organized by the 3’ closing base pair.

#### Bulges

The bulge length distribution obeys an approximate exponential distribution (Figure 6A) consistent with the destabilizing energy of a bulge increasing as a function of length. When the bulge loop is of length 1 nt, the nucleotide is enriched for A, and depleted for G and C, when compared to global nucleotide frequencies in this database (Figure 6B). The strongest deviation from the exponential fit is at length 6 nt, which is also enriched for bulges with a GA mismatch (Supplementary Figure S7A).

**Figure 6.**
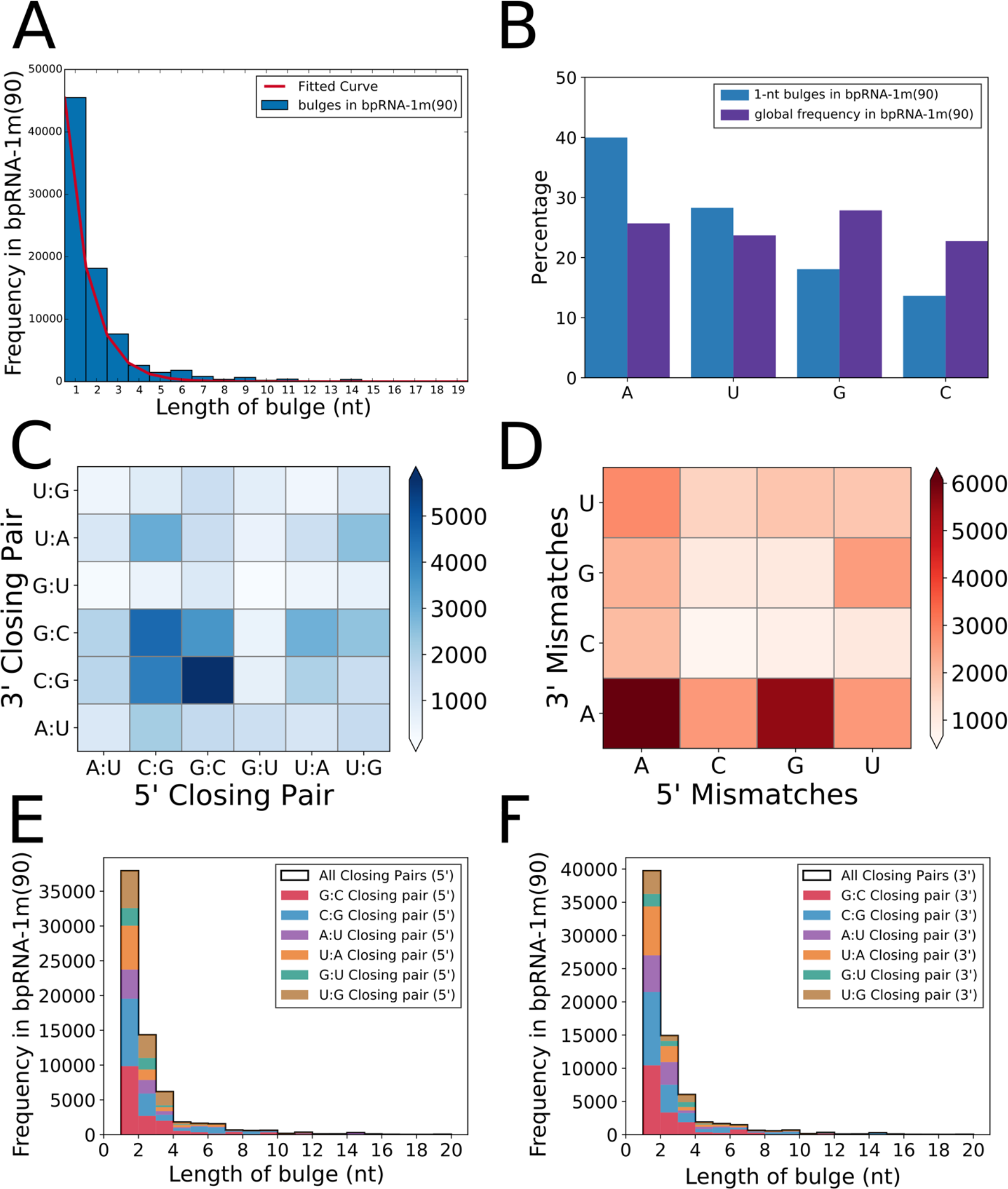
Bulges. **A.** Bulge length histogram. **B.** Nucleotide frequency in bulges of length 1. **C.** Heat map of closing base pairs. **D.** Heat map of mismatches. **E.** Bulge length distribution for different 5’ closing base pairs. **F.** Bulge length distribution for different 3’ closing base pairs.

Similar to internal loops, bulges show the highest enrichment for G:C at the 5’ and C:G at the 3’ closing pairs (Figure 6C). The majority of bulges are flanked by GC base pairs, but for bulge loops less than 3 nt other flanking base pairs are common (Supplementary Figure S7B-D). In addition, while GA mismatches are the most common for internal loops, the most common mismatch for bulges is AA, with GA the second most common (Figure 6D). The largest asymmetry between 5’ and 3’ closing base pairs was observed for U:G 5’ closing pairs for bulges less than 4 nt (Figure 6E,F). Internal loops and bulges show similar trends for lengths when binned by closing pairs, but with bulges having a more sharply decaying distribution.

#### Multiloops

Based on analysis of bpRNA-1m(90), we found that multiloops branches (junctions) of size 3, 4, and 5 are the most common and multiloops of greater than size 6 nt are very rare (Figure 7A). Additionally, the distributions of branch lengths for these common multiloop branch-counts indicates that multiloops with 4 branches are significantly enriched for multiloop branches of zero length (Figure 7B) which is found in “flush stacking”(17). This pattern is consistent with the fact that multiloops with four branches have more opportunities to be stabilized by coaxial stacking when the branches are zero length. In contrast, two helices in a multiloop with three zero-length branches would still be offset asymmetrically by the width of the third helix.

**Figure 7.**
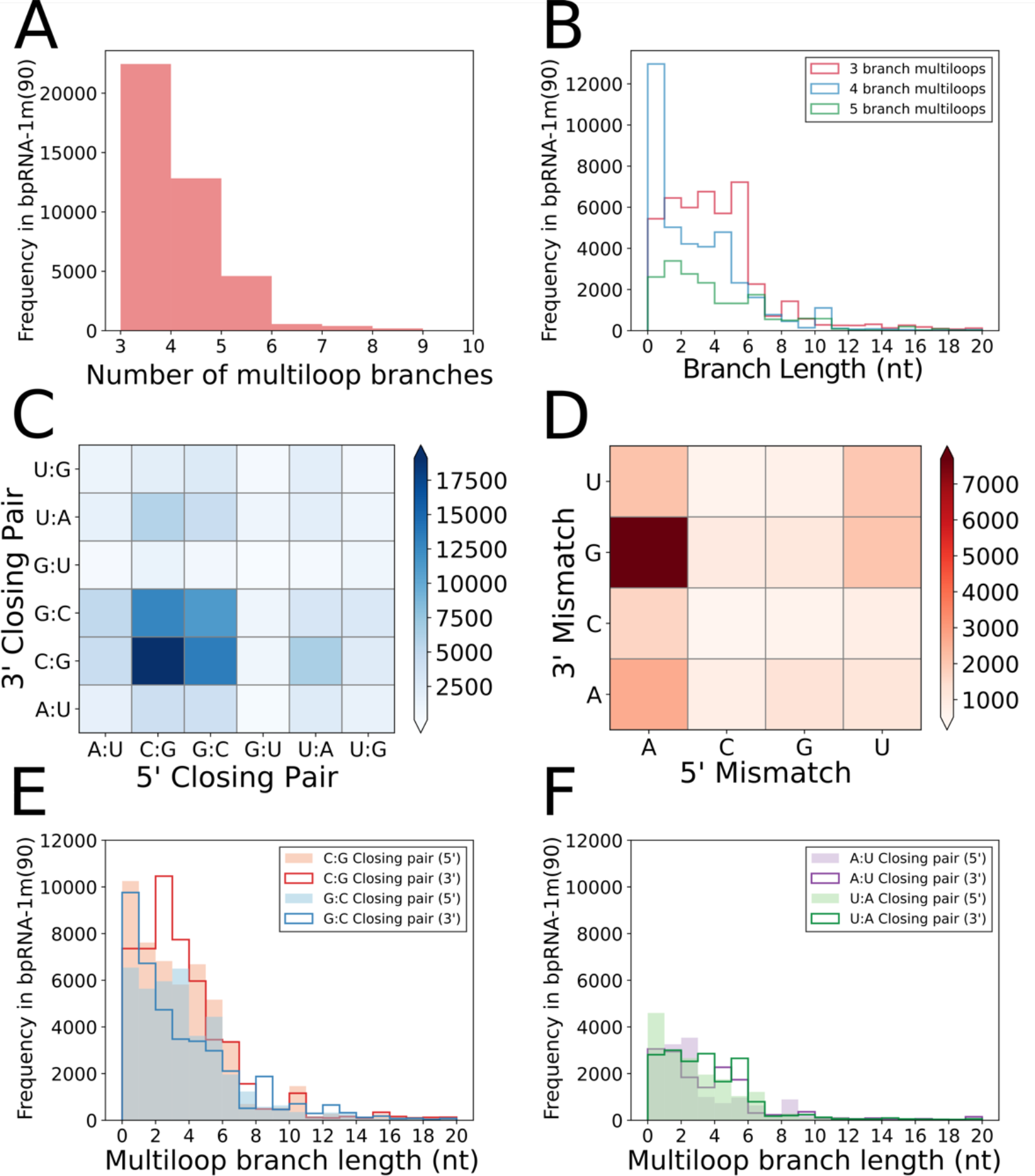
Multiloops. **A.** Histogram of branch number for all multiloops in bpRNA-1m(90). **B.** Branch length for multiloops with different branch numbers. **C.** Closing pair heat map. **D.** Mismatch heat map. **E.** Length distribution for different GC closing base pairs **F.** Length distribution for different AU closing base pairs.

Heat maps of the frequency of each closing base pair in multiloops branches demonstrates that most of the closing base pairs in multiloops tend to be C:G for 5’ closing pairs, and G:C for 3’ closing pairs—the opposite of internal loops (Figure 7C). This pattern for the closing pairs is the most common regardless of the number of branches (Supplementary Figure S8A-C). Overall, G:C and C:G closing pairs are significantly more common (Supplementary Figure S8D-I). In contrast to both internal loops and bulges, the most common mismatch pair for multiloops is AG (Figure 7D). Multiloop branches have a strong preference for GC-base pairing, with loops of length 0 showing a preference for C:G closing pairs, and loops of length 2 showing a preference for G:C closing pairs (Figure 7E-F).

#### Stems

Each stem in the database can be considered an instance of a “stem type”, such as CAG:CUG. To avoid double-counting, we alphabetically sort the two strands to form a distinct type. The full bpRNA-1m database contains of 2,075,928 stems that are instances (tokens) of 44,307 stem types, and bpRNA-1m(90) has 335,877 stems and 34,424 stem types. The frequency of stem type occurrences obeys a Zipfian distribution(48,49), as observed in Figure 8A. The frequency *f* of occurrence of stems follows the equation, *f* = *Ar*^−*s*^ where *r* is the rank of the stem when sorted by frequency, and the scale factor *s* ≈ 1.005, extremely close to the idea Zipf relationship of *s* = 1. The frequency of occurrence of stems does not correlate perfectly with the energy of the stem sequence, because longer stems are typically less frequent.

**Figure 8.**
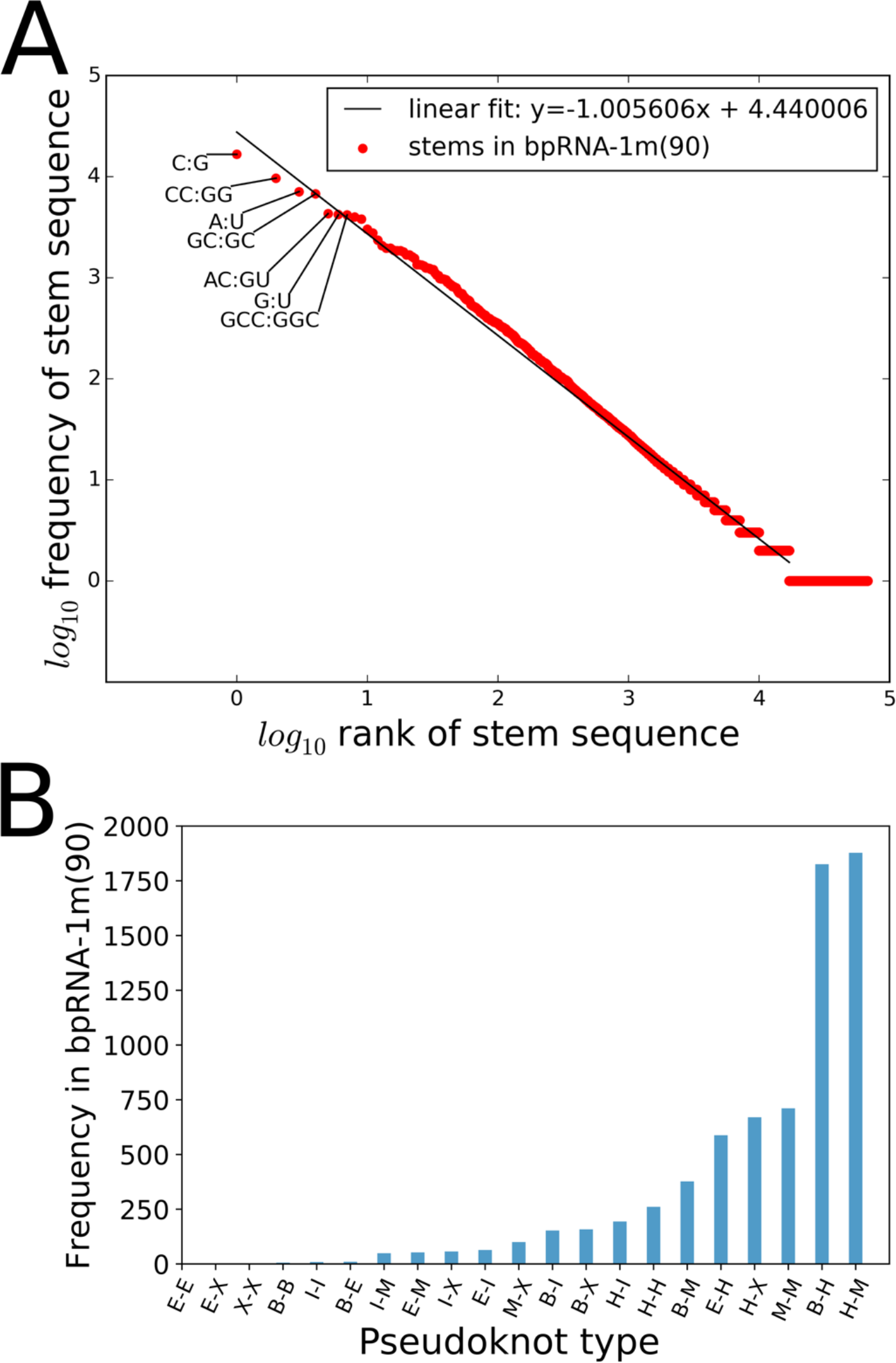
Stems and Pseudoknots. **A.**The frequency of stem types compared to their rank has a Zipfian distribution with a scale factor approximately equal to −1.00. **B.** bpRNA classifies pseudoknots by the loops that their base pairs connect when the pseudoknots are removed.

#### Pseudoknots

Around 12% (3,320) of RNA structures have at least one pseudoknot (PK) in bpRNA-1m(90) (Supplementary Table S2). Most PK-containing RNAs have only one PK; however, many RNA secondary structures contain more than one PK. Overall, there are 7,164 PKs in this data set. To get a sense of most frequent loop types forming the PK structures in our set, we plotted the frequency of each type of PK in Figure 8B. The most frequent type is between multiloops and hairpin loops, followed by bulges and hairpin loops. kissing hairpins (H-H), which are commonly studied(50,51), are the 7^th^ most common. Consistent with our expectations that dangling ends and external loops cannot form pseudoknots with each other because such an interaction would form a multiloop and not a PK, our annotations do not find any examples of this. Analyzing base pair information per pseudoknot structures suggests that PKs with three base pairs are the most frequent in our dataset and there are only four PKs in bpRNA-1m that have 12 base pairs, the largest observed in bpRNA-1m (only one PK with length 12 observed in bpRNA-1m(90)).

#### Non-Canonical Base Pairs

The C:G/G:C base pairs in both bpRNA-1m and bpRNA-1m(90) outnumber any other base pairs. In addition to Watson-Crick (base pair interaction between C and G or A and U) and wobble base pairs (G:U pairs), there are other nucleotide interactions observed in the databases we have compiled, commonly referred to as non-canonical base pairs. Even though the canonical base pairs (Watson-Crick and wobble pairs) are more common in RNA secondary structures formation, non-canonical base pairing is important in the formation of the tertiary structures. We observe 9.1% of the base pairs in bpRNA-1m(90) are non-canonical. In 44.8% of these non-canonical pairs occur in the middle of a stem surrounding by canonical pairings, whereas only 7.2% are isolated base pairs. Also, about 1.4% of these special pairings are involved in pseudoknot formation (All stats are based on bpRNA-1m(90)). Table 4 shows the frequency of each type of base pairs in bpRNA-1m and bpRNA-1m(90). A:G/G:A, and A:C/C:A are the most common non-canonical pairs in both bpRNA-1m and bpRNA-1m(90), and C:C are the least frequent.

**Table 4.** Number of canonical and non-canonical base pairs in bpRNA-1m and bpRNA-1m(90)

| Base pair | bpRNA-1m | bpRNA-1m(90) |
| --- | --- | --- |
| C:G | 5,027,894 | 747,110 |
| A:U | 2,232,052 | 410,641 |
| G:U | 1,137,821 | 174,545 |
| A:G | 239,066 | 32,564 |
| A:C | 105,964 | 26,074 |
| U:U | 87,396 | 18,958 |
| C:U | 56,063 | 18,587 |
| G:G | 51,959 | 11,820 |
| A:A | 39,072 | 10,499 |
| C:C | 21,421 | 7,748 |

### Summary and Conclusions

We have developed the bpRNA annotation approach to reliably produce intuitive secondary structure annotations from base pairing data to help with understanding RNA structure. Our efforts to provide annotations that are more informative and generally applicable than previous approaches have yielded many new strategies for representing RNA structural data such as the structure array, which makes the structure easier to read and visualize by providing a character label for each nucleotide of the dot-bracket representation. Likewise, the structure type file represents a detailed annotation, covering each nucleotide of the sequence. Separating the structure into segments—base paired regions interrupted by only bulges and internal loops—provides facile identification of multiloops and external loops, even when their length is zero. bpRNA also creates accurate dot-bracket representations for both simple and complex pseudoknot-containing RNA secondary structures.

We applied bpRNA to create a large integrated meta-database of single molecule RNA secondary structure that we have assembled from seven different sources (bpRNA-1m). With this large meta-database and the RNA structural information that bpRNA provides, there is an opportunity for a number of applications. The annotations produced from bpRNA could be used to improve the source databases used to build bpRNA-1m. Expanded structure annotations could enable the calculation of a next generation of thermodynamic parameters. The data set generated by bpRNA is large enough to enable training and testing machine learning algorithms for the prediction of RNA structure. Moreover, by restricting to only include single molecule structures, this dataset can serve as a benchmark for RNA secondary structure prediction algorithms, which typically take a single sequence as input.

We have used the annotation details and structural features produced by bpRNA to identify several statistics trends in bpRNA-1m(90), which contains over 28,000 sequences that are less than 90% similar, over 10 times the size of previous similar refined data(52). While some of these trends represent patterns of thermodynamic stability, future studies are needed to expand this analysis with more structures, or judiciously filter the data for a more refined structural analysis.

## DATA AVAILABLITY

All data and scripts are accessible to download in http://bprna.cgrb.oregonstate.edu/index.html

## SUPPLEMENTARY DATA

Supplementary Data are available at NAR Online

## AKNOWLEDEGEMENT

The authors would like to thank Professors P. Andy Karplus and David H. Matthews for input and feedback that improved the manuscript.

